# Supra-resonant wingbeats in insects

**DOI:** 10.1101/2025.05.07.652680

**Authors:** Ethan S. Wold, Rundong Yang, James Lynch, Ellen Liu, Nick Gravish, Simon Sponberg

## Abstract

Powering small-scale flapping flight is challenging, yet insects sustain exceptionally fast wingbeats with ease. Since insects act as tiny biomechanical resonators, tuning their wingbeat frequency to the resonant frequency of their springy thorax and wings could make them more efficient fliers. But operating at resonance poses control problems and potentially constrains wingbeat frequencies within and across species. Resonance may be particularly limiting for the many orders of insects that power flight with specialized muscles that activate in response to mechanical stretch. Here, we test whether insects operate at their resonant frequency. First, we extensively characterize bumblebees and find that they surprisingly flap well above their resonant frequency via interactions between stretch-activation and mechanical resonance. Modeling and robophysical experiments then show that resonance is actually a lower bound for rapid wingbeats in most insects because muscles only pull, not push. Supra-resonance emerges as a general principle of high-frequency flight across five orders of insects from moths to flies.

---

Among the four evolutionary lineages of flying organisms, insects uniquely produce rapid, powerful wingbeats at the centimeter scale from the low whir of a giant silkmoth to the near-kHz hum of a biting midge (Fig. 1a) (*1, 2*). Wingbeat frequency is a critical determinant of aerodynamic power production (*3–6*), but is only weakly predicted by body size at the species level. For example, bumblebees flap at 180 Hz (*7*), but bee-mimicking hawkmoths, *Hemaris diffinis* (*8*), flap at 60 Hz despite being nearly the same size. Resonance is a popular explanation for this many-to-one mapping between body size and wingbeat frequency. Most insects fly by deforming an elastic exoskeleton with ultrafast flight muscles (*9–15*). Flapping at their resonant frequency theoretically allows for the costs of rapid wing acceleration to be offset by elastic energy storage, but at the expense of frequency modulation capacity. While wing-clipping experiments (*3, 16*) and models (*9, 17*) point to insects being resonant, it is an unresolved question whether insects flap at resonance. Slow-flapping (*<*100 Hz), synchronous insects like moths may not be overly restricted by resonance because their wingbeats are paced by time-periodic neural signals, which can match or exceed the resonant frequency. However, resonance may be particularly constraining for fast-flapping (typically *>*100 Hz) asynchronous insects that generate self-excited wingbeats with specialized muscles that activate in response to mechanical stretch (*18–20*) (Fig. 1b). By combining new measurements of insect muscle and exoskeleton with models of ‘spring-wing’ dynamics, we investigate whether insects flap at resonance across taxa and flight mode, and if not, what timescales set their wingbeat frequencies.

**Figure 1:**
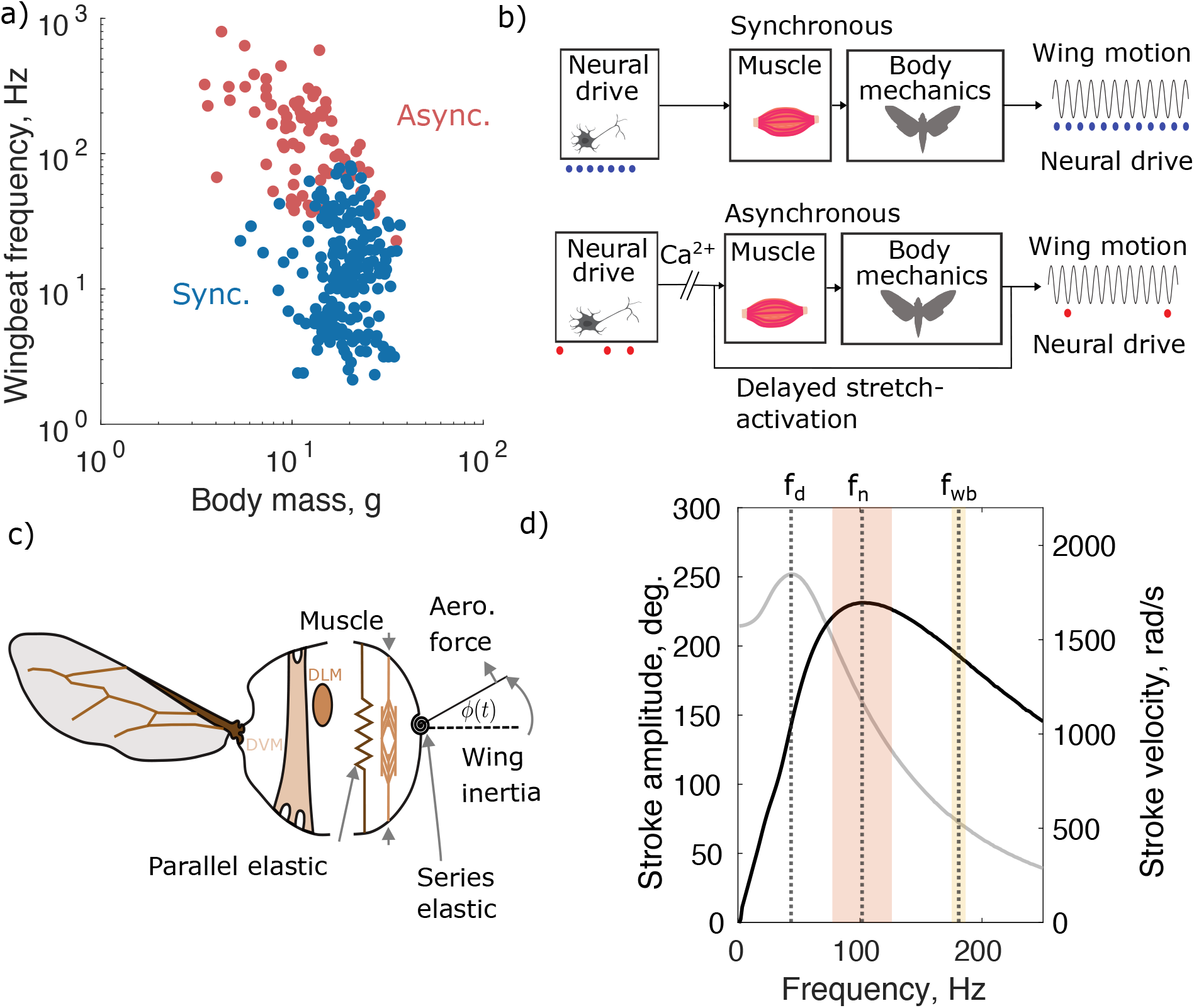
a). The division between synchronous and asynchronous insects helps explain the wide variation in insect wingbeat frequency. Data replotted from (*1, 2*). b) Insects with synchronous muscle produce wingbeats at a frequency set by the neural drive to the flight muscles (blue dots). Insects with asynchronous muscle produce faster wingbeats that are decoupled in frequency from the underlying neural drive (red dots). c) Schematic of a bumblebee showing flight muscle anatomy on the left, and a biomechanical model on the right. d) Simulated displacement resonance curve (grey) and velocity resonance curve (black) of a bumblebee, assuming a sinusoidal forcing. Displacement resonant frequency (*f*_*d*_) and velocity resonant frequency (*f*_*n*_) are both below wingbeat frequency (*f*_*wb*_). Orange and yellow bars show 95% confidence intervals of the mean for velocity resonance and wingbeat frequencies respectively.

## Bumblebees flap above their resonance frequency

First, we focus our attention on the bumblebee, whose flight kinematics, morphology, and behavior have been characterized in detail. Using materials testing in the context of a ‘spring-wing’ model of an insect’s elastic thorax, and the inertial and aerodynamic forces acting on the wing (*8, 10, 11, 20*), we demonstrate that asynchronous bumblebees flap above their resonant frequency (Fig. 1c-d). We estimated the resonance frequency of *Bombus impatiens*, by combining measurements of bulk thoracic stiffness with estimates of wing inertia and wing-hinge transmission ratio. The bumblebee’s undamped resonant frequency (*f*_*n*_) is a function of the measured thorax stiffness (*k*), the wing hinge transmission ratio (*T*, the ratio of angular wing displacement to muscle displacement with units rad m^− 1^), and the inertia of the wings and added mass of air around the wing (*I*, see SI for extended description of all parameters),

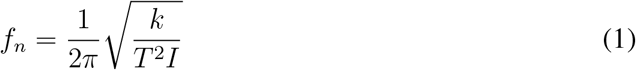

The elastic thorax and main flight power muscles (Fig. 1c) are in a parallel configuration that drive indirect actuation of the wing. We ignore series elasticity of the wing hinge, which is likely small and would widen but not alter the location of the undamped resonant frequency peak (*13*) (see Supplementary Discussion). We measured the isolated thorax stiffness of bumblebees using vibrational testing and found it to be 4.1 kN/m (see SI section 2.1) Setting all parameters from empirical measurements, we arrive at an undamped resonant frequency of *f*_*n*_ = 94.9 Hz, which is 44% lower than average wingbeat frequencies (180 Hz) (*7*) (Fig. 1d). Our measured thorax elasticity does not take into account active stiffness contributed by the flight muscles. Since the upstroke and downstroke muscles are antagonistic, when one contracts the other is stretched under near-tetanic activation. We estimate, conservatively, that active muscle stiffness is equal to the summed stiffness of both pairs of flight muscles, increasing the total thoracic stiffness to 6.4 kN*/*m and the resonant frequency to 102 Hz (*21*) when considering stiffness contributions from exoskeleton and active muscle. Propagating error in the three parameters *k, T*, and *I*, we find that supra-resonant bumblebee wingbeats are robust to reasonable measurement error in thorax or wing properties, resulting in resonant frequencies ranging from 76 to 124 Hz (Fig. 1d).

We can also reverse the analysis to ask what thorax stiffness is necessary for wingbeats to be at the resonance frequency. This would require a stiffness of 21 kN*/*m, which is beyond the largest stiffness that we measured and far exceeds any comparable stiffness measurements (≈ 2 kN/m) (*15, 22*). Incorporating the effects of aerodynamic force production or internal thoracic damping into the resonance calculation can only further depress the estimated resonant frequency below wingbeat frequency (i.e. damped displacement resonance) (*8, 11*). More complex resonant models could create nonlinear resonance at higher harmonics, but we are interested here in the fundamental resonance from the exchange of inertial and elastic energy during the wingstroke. Thus, in the absence of evidence that substantial elasticity is missing from our measurements, we conclude that bumblebees are supra-resonant.

## Stretch-activated dynamics of asynchronous muscle

The discrepancy between bumblebee resonant and wingbeat frequencies motivated us to consider how the physiological process of stretch-activation in muscle can enable supra-resonant flight in asynchronous insects. Asynchronous muscle generates active force in response to a rapid stretch. This mechanical stretch-activation was measured previously by stretch-hold experiments (*18–20, 23*) (Fig. 2a-b). The force response of isolated insect flight muscle to a step length change under tetanic activation has a shape that is composed of four phases (*24*) (Fig. 2c). The first two phases are fast, associated with the viscoelastic response of the muscle tissue. The slower third and fourth phases comprise the delayed stretch activation (dSA) force, which can be described with a single characteristic timescale, *t*_*o*_ (the time taken to achieve peak dSA force after the end of stretch (*20*)), and a constant, *ε*, the ratio of the rates of force decay (*r*_4_) to force rise (*r*_3_) (Fig. 2d). An analogous process, delayed shortening de-activation (dSD) occurs following rapid shortening (Fig. 2e, g) and is the inverse of dSA. We hypothesized that an asynchronous insect can flap above resonance if its *t*_*o*_ is sufficiently fast with respect to its natural period, the reciprocal of natural frequency 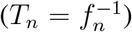. In this case the resulting wingbeat frequency is set not just by the resonant mechanics, but by a combination of muscular (*t*_*o*_) and mechanical (*T*_*n*_) timescales.

**Figure 2:**
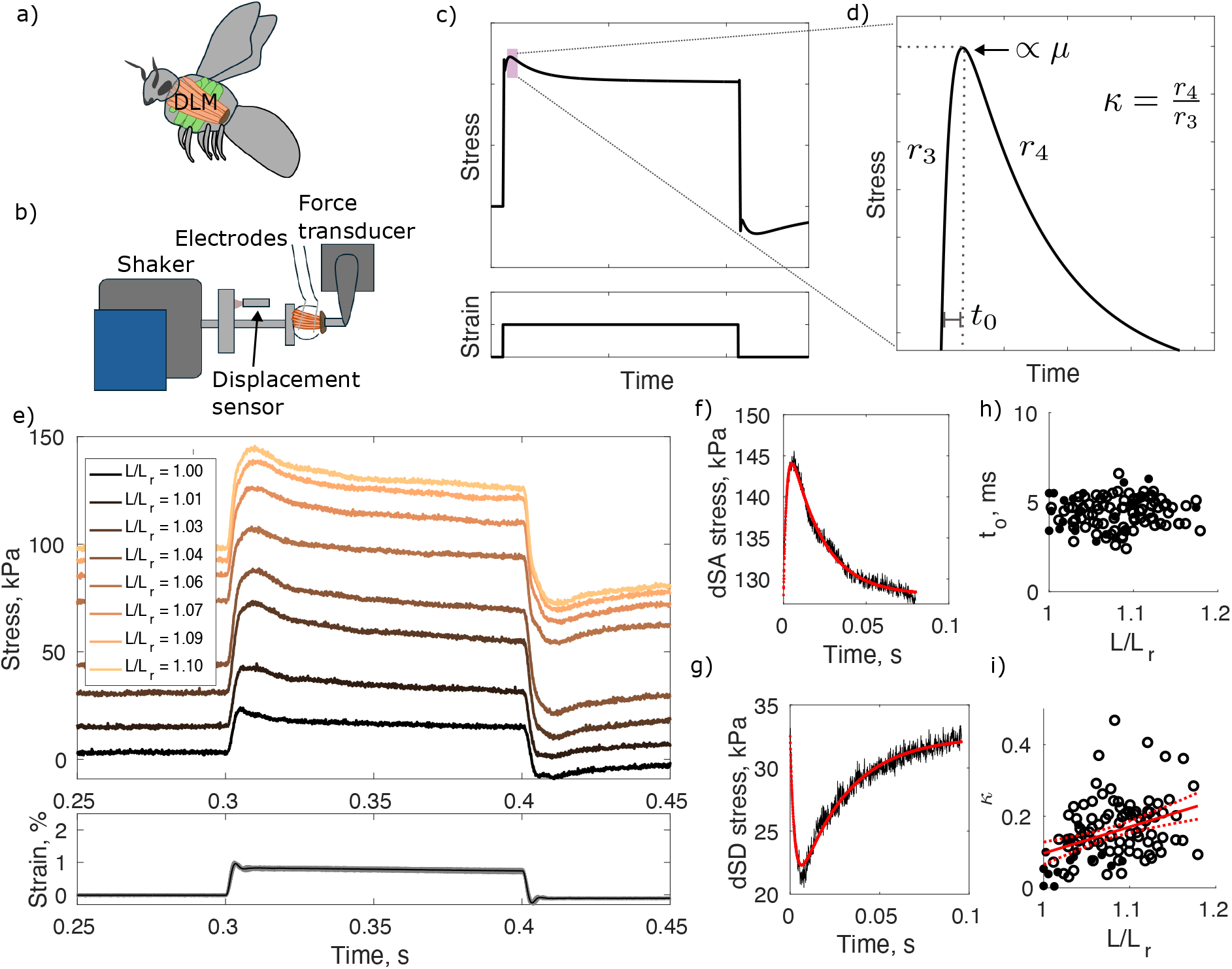
a). Location of the DLM (downstroke) muscles in a bumblebee. b). Schematic of muscle physiology apparatus used to apply step strains along the line-of-action of the DLM under tetanic stimulation. c). Zoomed out cartoon of a single stretch-hold-release-hold trial with purple bar blown up in panel d) Phases three and four of the dSA response, with corresponding rates of force rise (*r*_3_) and force decay (*r*_4_) notated. *t*_*o*_ is the time until peak dSA force is reached, and κ is the ratio of *r*_4_ to *r*_3_. *µ* is proportional to the height of the dSA response. e) dSA characterization experiments from a single individual. A 1% stretch was applied under tetanic stimulation at multiple starting lengths. f) dSA phases 3 and 4 from a single trial from a single individual. Red line denotes a double-exponential fit. g). dSD phases 3 and 4 from a single trial from a single individual. h) *t*_*o*_ computed across all individuals at all starting lengths (n=10 individuals). i) κ computed across all individuals at all starting lengths. Closed and open circles correspond to dSA and dSD measurements respectively. Red line shows a linear regression with 95% confidence intervals.

To measure a bumblebee’s dSA timescale under typical flight conditions, we had to conduct new stretch-hold experiments on isolated *B. impatiens* DLMs at the a realistic flight temperature of 40^°^C (*25, 26*) (Fig. 2a-e). While some measurements of bumblebee stretch-activation exist, we are the first to make them in intact whole muscle at a realistic body temperature. We measured the bumblebee stretch-activation timescale, *t*_*o*_, to be 4.4 *±* 1.0 ms, nearly the duration of an entire wingbeat, which did not change depending on muscle length at the onset of stretch (Fig. 2f-h). This value of *t*_*o*_ is somewhat faster than the only comparable characterizations in bumblebees at lower temperatures (≈ 5 ms) (*21, 27*), which may be because dSA rate kinetics are known to speed up with temperature (*23*). It is also substantially slower than the ≈ 2.5 ms necessary for dSA alone to drive 180 Hz wingbeats in response to stretch (the duration of downstroke after being stretched during upstroke). The ratio of rates, *ε*, was weakly correlated with resting muscle length and on average equal to 0.17 *±* 0.03 (Fig. 2i). Thus, the muscular stretch response is faster than the timescale of mechanical resonance.

## Asynchronous insects flap at or above resonance

To test our hypothesis that a sufficiently fast stretch-activation timescale (i.e. *t*_*o*_) can enable supra-resonant wingbeats, we developed a biophysical model of asynchronous muscle-driven resonant flight. We drive a ‘spring-wing’ model of the insect’s thorax and wings (*8, 10 11*) with a simplified description of stretch-activated muscle, rooted in our stretch-hold experiments in bumblebees (*20, 30*). Thus, we capture the essential dynamics of both stretch-activation and resonant mechanics.

The model consists of two coupled, second-order differential equations. The oscillatory dynamics of the wingstroke angle, *ϕ*(*t*) are parameterized by *k*_*l*_, *T, I*, and an aerodynamic damping coefficient Γ (Eq. 2). The stretch feedback-driven muscle forcing (Eq. 3) is parameterized by *t*_*o*_ and κ (see Supplementary Methods section 1.6.2). There is one free parameter, *µ*, which can be set by matching the amplitude of the output wingbeats to free-flight bumblebee wingstroke amplitude. *µ* represents the strength of the dSA forcing and makes the dSA force *in-vivo* proportional to the height of the two-phase response of the muscle to rapid stretch (Fig. 2c-d). Because the forcing is entirely state dependent and not prescribed exogenously (e.g. by the nervous system) the system oscillates at an emergent frequency (See Supplementary Results 2.2).

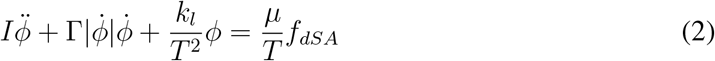

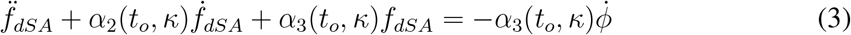

Simulating Eqs. 2-3 and evaluating the resulting *ϕ* (*t*) wing stroke trajectories over a wide range of muscular and mechanical timescales (*t*_*o*_ and *T*_*n*_) reveal that resonance is a lower bound on emergent wingbeat frequency (Fig. 3a). There exists a large region of parameter space in which asynchronous insects can flap significantly above resonance. Consistent with observations from modifying the wing inertia of insects (*3,16,31*), the resonance frequency of the insect increases as the emergent flapping frequency goes up, regardless of *t*_*o*_ (Fig. 3a). However, this does not mean that the flapping frequency is at resonance. Fast stretch-activation, low *t*_*o*_, results in supra-resonant wingbeats, while slower *t*_*o*_ result in wingbeat frequencies that collapse onto the wingbeat-resonant frequency equivalency line. Thus, stretch-activated resonant systems are not constrained to resonance, and can oscillate supra-resonantly with the right combination of *t*_*o*_ and *T*_*n*_. Using our measured flight muscle stretch-activation timescale, we estimate a bumble-bee wingbeat frequency which exceeds resonance by 33%, in agreement with our experimental characterization of supra-resonance using thorax materials testing.

**Figure 3:**
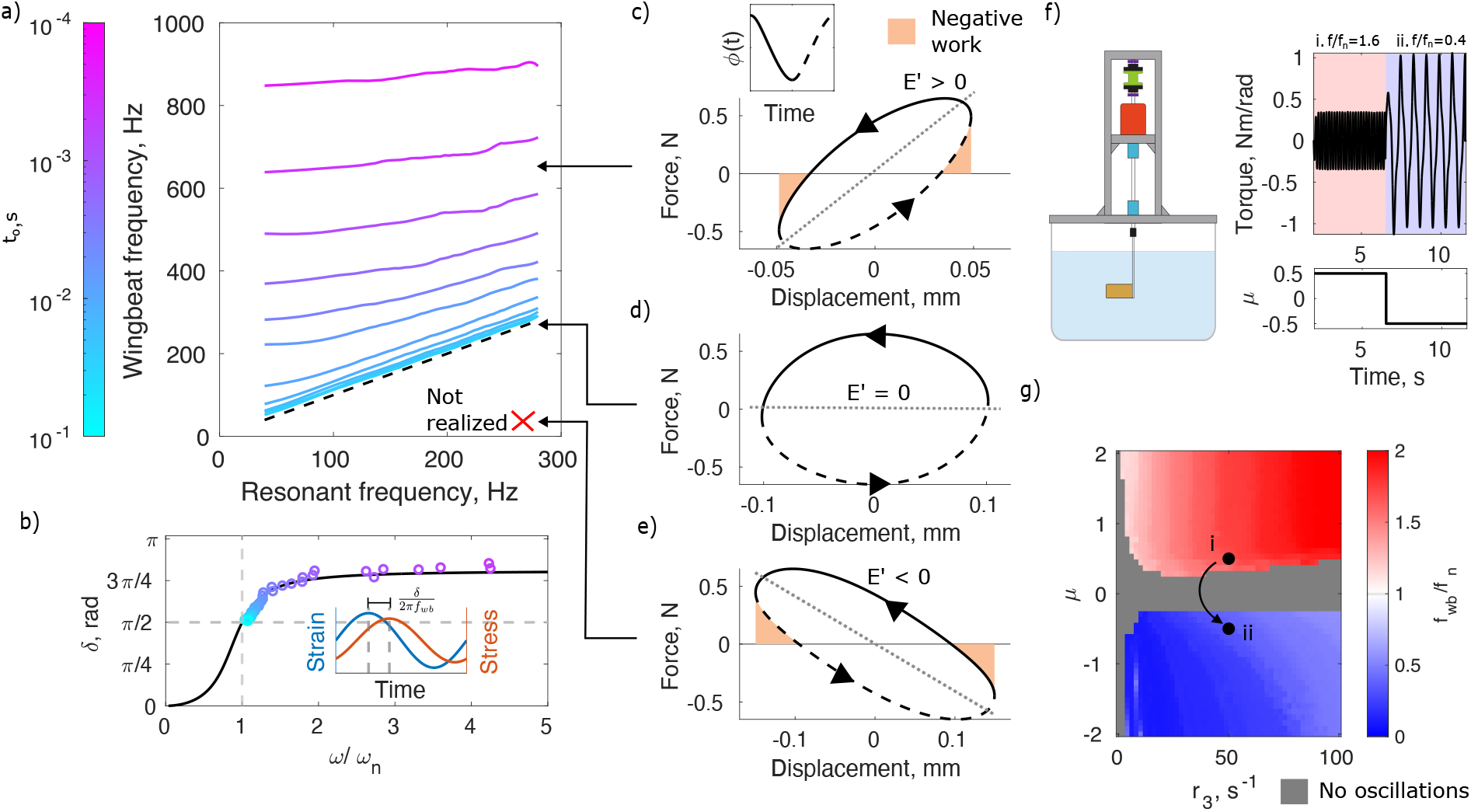
a). Parameter space of emergent wingbeats over a wide set of *t*_*o*_ and *T*_*n*_ that encompasses the physiological range for most insects. For any fixed value of *t*_*o*_, there exists a linear relationship between resonant and emergent frequencies. At large *t*_*o*_, this linear relationship collapses onto the equivalency line. As *t*_*o*_ decreases, wingbeat frequencies become supra-resonant, and increasingly independent of resonant frequency. Every point in simulation has been normalized such that steady-state peak-to-peak stroke amplitudes match bumblebee *in-vivo* conditions. b). Phase lag between stress and strain as a function of normalized frequency. Black line shows phase for a synchronous insect and colored dots show predictions from the asynchronous model for varying *t*_*o*_, with the same color scale as in panel a). c-e). Work loop in force-displacement space for operation above (c), at (d), and below (e) resonance. Dotted lines correspond to upstroke and solid lines correspond to downstroke. At resonance, no negative work is required of the flight muscle. Above and below resonance, negative work is done in the second half and first half of each half-stroke, respectively. f). Robophysical experiment where the sign of the dSA force is changed from positive to negative, causing oscillations to transition from supra-resonant to sub-resonant. g). Space of asynchronous wingbeats in a robophysical flapper, demonstrating that transitions between sub- and supra-resonance occur only when the dSA force sign changes (i.e. across the *µ* = 0 boundary). Parameter values corresponding to the transition in f). are shown by the arrow from i. to ii.

## Muscle’s asymmetry enforces supra-resonance

Why are asynchronous flappers apparently only able to flap at or above resonance? This constraint emerges from muscle producing a positive force in response to stretch (i.e. always ‘pulling’ and never ‘pushing’), such that increases in muscle stress will always lag positive muscle strain (muscle extension) (Fig. 3b, inset). This phase lag is proportional to the stretch-activation timescale (*t*_*o*_), but can never be below a quarter cycle (π */*2 radians) because increasing *t*_*o*_ results in a frequency that asymptotes to resonance. As *t*_*o*_ increases, δ approaches *π /*2; as *t*_*o*_ decreases, *δ* approaches a maximum. Thus, stretch-activated wingbeats can never be sub-resonant, which would require δ *< π /*2 (Fig. 3b, colored points). This contrasts with a synchronous insect that has neurally activated muscle, which can theoretically achieve 0 *< δ < π* by modulating the timing of neural signals to the flight muscles (Fig. 3b, black line). This would be equivalent to activating the muscle such that it produces significant force in the half cycle prior to lengthening (*32–36*).

The necessity of asynchronous insects to be supra-resonant can be visualized in the work-loop representation of muscle function, which visualizes muscle work output as the area enclosed in active force vs. displacement space (*37–40*). In this space, the effective storage modulus, *E* ′, of the workloop represents the elastic component of the total muscle force necessary to supply all energy requirements for flight (see Supplementary Results 2.3). *E*^′^ = 0 represents perfect exchange between wing kinetic energy and thoracic elastic energy (i.e. *f*_*wb*_ = *f*_*n*_) (Fig. 3d). *E*^′^ ≠ 0 represents non-resonant conditions where the thorax is either too stiff or not stiff enough with respect to the wingbeat frequency (Fig. 3c,e). The hysteresis of the loop is proportional to the phase lag δ between force and displacement. The resulting area within the loop is the effective loss modulus, *E*^′ ′^, representing the mechancial energy generated by the muscle.

At steady-state, the muscle forcing *f*_*dSA*_(*t*) will always follow a counter-clockwise ellipse (negative loss modulus or positive net work) with a nonnegative storage modulus. This enforces a phase lag, *δ*, of *π/*2 *< δ < π* in stress with respect to strain. If the insect is supra-resonant, its muscle will produce negative work, dissipating energy, during the second half of each halfstroke while the wing is decelerating (Fig. 3c, shaded area). Muscle assisting in the slowing of the wing on each half cycle is only possible in a supra-resonant system, since the thoracic spring will not be stiff enough to absorb all of the wing’s kinetic energy before it reaches its extremal positions. At resonance, the storage modulus of the muscle is identically zero and no negative work is required at any point in the wingbeat (Fig. 3d). Below resonance, an overly stiff thoracic spring would cause rapid wing acceleration that is counteracted by muscle dissipating energy in the acceleration phase of each half-stroke (Fig. 3e, shaded area). Thus, a sub-resonant asynchronous insect would have to generate negative work directly following stretch, which is incompatible with the polarity of muscular dSA force response. Muscles only pull, they do not push.

## Supra- and sub-resonance realized through a robophysical model

While the physiological limitations of biological muscle limit oscillations to at or above resonance, engineered actuators are not necessarily limited to this regime. In contrast, they can push and pull. Following from the sub-resonant work loop (Fig. 3e), we predict that an engineered system with muscle dSA-like actuators that pushed instead of pulled should be able to realize sub-resonant asynchronous wingbeats. We demonstrate asynchronous sub-resonant oscillations in a dynamically scaled robophysical flapping wing by controlling a brushless DC motor with a velocity feedback-driven forcing analogous to dSA (Eqs. 2-3) (*30*). This system is also unable to oscillate below its resonant frequency, when controlled with a muscle-like dSA forcing (Fig. 3f, i.). However, by changing the sign of the dSA force such that a negative (i.e. pushing) force follows stretch, stable sub-resonant oscillations emerge that are bounded above by the resonance frequency (Fig. 3f, ii.). By systematically changing *µ* in the model that controls our roboflapper motor, we see the boundary of switching between supra- and sub-resonant behavior is at *µ* = 0, where the sign of the dSA force flips (Fig. 3g). Thus, we demonstrate that the physiology and arrangement of antagonistic stretch-activated muscles in asynchronous insect thoraces constrain them to supra-resonant wingbeats.

Sub-resonance is realizable in some biological muscle-driven systems as well. For instance, unlike asynchronous insects, synchronous insects can, in theory, be sub-resonant. They could neurally activate their muscles at timings that would enable negative work production in the first part of each half-stroke. Practically, this would require the downstroke muscle (or a combination of muscles) to produce significant force during the beginning of upstroke, and vice versa. This would require either multiple downstroke or upstroke muscles or a large degree of coactivation which would likely reduce the production of useful work from the muscles. While we do not see such activation patterns in insects, they are common in terrestrial locomotion especially where multiple muscles operate in synergy at a joint or in a limb (*33–36, 41*). Indeed, some terrestrial animals like kangaroos are sub-resonant (*42*), but do not have to contend with the asynchronous muscle dynamics present in bumblebees or our robophysical flapper.

Supra- and sub-resonant systems also exhibit a key difference in how they modulate power output outside of the steady state. In a supra-resonant system, acceleration is muscle-driven and spring-assisted, with negative muscle work (dissipation) coinciding with the deceleration phase of the wingstroke (Fig. 3c). The muscle assists the spring. This enables supra-resonant systems to inject additional accelerative power via transient increases in agonist muscle force. However, in sub-resonant systems, acceleration is driven by the spring and braked by the agonist muscle (Fig. 3e). Positive power production is limited by the spring’s capacity to return elastic energy and additional agonist muscle force would only decelerate the wing faster. Thus, an important benefit of supra-resonance for asynchronous insects may be to maintain the capacity to transiently boost wing acceleration via positive muscle power production in the first part of each half-stroke (Fig. 3c).

## Two timescales pace asynchronous wingbeats

Having validated our model in bumblebees, we used recent characterizations of *Drosophila melanogaster* muscle stiffness and flight mechanics to show that this widely-studied fruitfly is also supra-resonant, although to a lesser degree than *B. impatiens. D. melanogaster* has more than three orders of magnitude less mass than a bumblebee, but paradoxically has a wingbeat frequency of around 200 Hz, which is very close to that of *B. impatiens* (*43*). Measurements of fruitfly delayed stretch activation timescale, *t*_0_, range from 5-8 ms which is slower than our measured *t*_*o*_ for bumblebees (Fig. 2h) (*44*). We hypothesize that fruitflies have evolved a similar wingbeat frequency to bees despite being much smaller in part by having relatively slow stretch-activation in comparison to their natural period. Indeed, fruitfly wing inertia is roughly four orders of magnitude smaller than that of a bumblebee, which in isolation would suggest a resonant frequency far greater than ≈ 100 Hz we measured in bees.

To test this hypothesis, we first need an estimate of the bulk stiffness of the fruitfly thorax. While exoskeletal stiffness values for fruitflies have not been measured, functional stiffness in the fly thorax is thought to be dominated by active muscular elasticity, rather than parallel thoracic stiffness. This is due to fruitflies’ combination of small wing inertia compared to bees, and asynchronous muscle which has much higher resting stiffness than synchronous muscle (*19, 21 28, 45*). We quantify whether existing estimates of muscle elasticity alone are sufficient to estimate resonance frequency, by deriving a new metric describing the contribution of muscular elasticity to bulk thorax stiffness: the ratio of the active muscle stiffness *k*_*muscle*_ to the stiffness that would be required to drive perfectly resonant wingbeats, *k* ^*\**^ = (2*πf*_*wb*_)^2^*T* ^2^*I* (Fig. 4a). This expression depends on nonlinear interactions between morphological (wing inertia, *I*), kinematic (wingbeat frequency, *f*_*wb*_), and biomechanical (transmission ratio, *T*) parameters. It is not a simple function of body size. Bumblebees and moth thoraces are dominated by exoskeletal stiffness, which is in excess of muscular stiffness by at least three-fold yet their free flight frequencies are still above their undamped resonance. In contrast, *Drosophila* muscle does supply nearly all the elasticity needed to drive wingbeats close to resonance (Fig. 4a).

**Figure 4:**
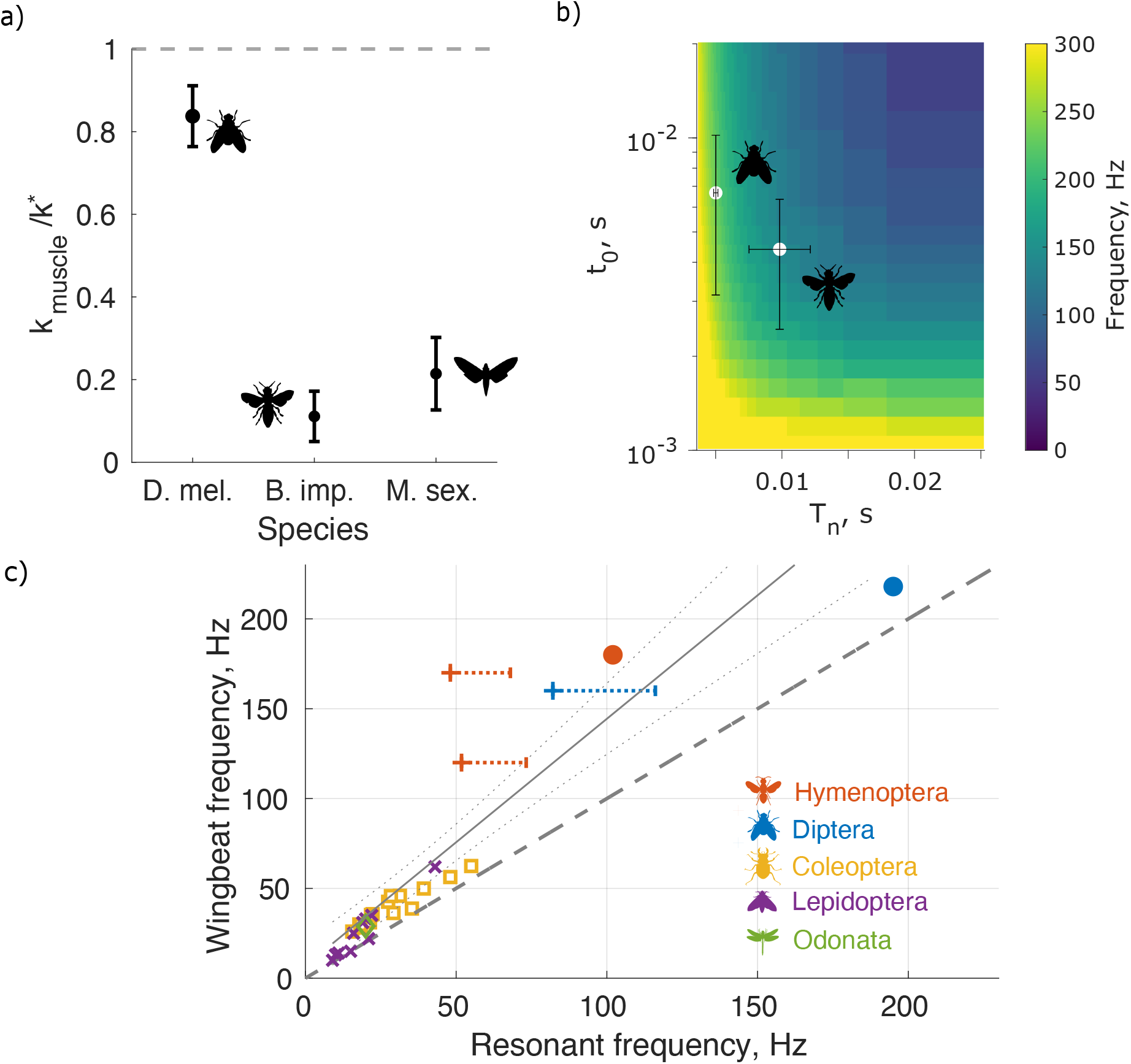
a). Fraction of perfectly resonant stiffness contributed by muscle alone for three insect species: fruitfly, bumblebee, and hawkmoth (*21, 28 29*). b). Emergent asynchronous frequency space as a function of muscle time constant (*t*_*o*_) and natural period (*T*_*n*_). Both fruitfly and bumblebee achieve similar wingbeat frequencies with different combinations of *t*_*o*_ and *T*_*n*_. c). Supra-resonant wingbeats across insects, compiling estimates from five insect orders. Markers - “× “(*8*), “+” (*22*), square (*18*), and diamond (*17*) - represent data from different studies. Dots show data from the current study. Dotted line shows equivalence of wingbeat and resonant frequencies. Orange bars show resonant frequency ranges assuming exoskeleton stiffness underestimates total thorax stiffness (exoskeleton + muscle) by up to a factor of two. Solid grey line shows a linear regression through all points, with dotted grey 95% CI.

Armed with a stiffness estimate for *Drosophila* we can then apply the same approach we took with bumblebees to test if they operate at their resonant frequency. When we estimate *Drosophila* resonant and wingbeat frequencies using Eqs. 1-3, we find a predicted emergent wingbeat frequency (221 Hz) that is supra-resonant at 18% in excess of resonance (187 Hz) (Fig. 4b). Supra-resonance arises from a combination of a much larger transmission ratio (ratio of wingbeat angle change to muscle displacement) by virtue of small body size as well as slower stretch-activation (longer *t*_*o*_). These frequencies are within the range of measured free-flight wingbeat frequencies. While series-elastic effects in moths and bees are not large enough to significantly impact our results (see Supplementary Discussion), high series-elastic compliance in *Drosophila*-scale flies may widen their resonance curves to the point where they can still achieve near-maximal resonant energy return while being supra-resonant (*13*). Thus, our modeling demonstrates how dSA causes a single frequency that exceeds *f*_*n*_ to emerge from a band of potentially equally efficient frequencies in insects with significant series compliance.We find that bees and flies realize similar asynchronous wingbeat frequencies through different combinations of *t*_*o*_ and *T*_*n*_ while remaining slightly (flies) to significantly (bees) supra-resonant (Fig. 4b).

## Supra-resonance as a general principle of insect flight

Our experimental and theoretical characterizations of resonance in bees and flies point to supra-resonance as a general principle of insect flight, but how broadly does it apply? Collating our results with the only other comparable characterizations of resonance (*17, 18 22*), we show that supra-resonance applies to asynchronous bees, flies, and beetles, and synchronous moths. Even the dragonfly, a synchronous insect with a direct flight muscle architecture appears to operate above its resonance peak. This suggests that supra-resonance is not limited to insects with indirect flight muscles so long as there is some degree of elasticity in the muscles or wing hinge (*17*). Thus, our results demonstrate a general pattern of supra-resonant wingbeats in insects, with all insects included flapping at or above their resonant frequency. Insects generally fall on a line with slope *>* 1 (slope= 1.32, *p <* 0.001, *r*^2^ = 0.77) but an intercept statistically indistinguishable from 0. Thus across taxa, insects maintain a roughly constant ratio of wingbeat to resonant frequency, with slower-flapping insects flapping extremely close to, but not below, resonance.

Interestingly, *Drosophila* lies closer to resonance than predicted by a line of best fit through all other species (slope= 1.84, *p <* 0.001, *r*^2^ = 0.80), suggestive that higher frequency insects may not continue to experience a divergence between resonant and wingbeat frequencies. This makes sense in the context of its larger *t*_*o*_ compared to that of a bumblebee (*44*), despite being orders of magnitude less massive. In addition, smaller insects achieve roughly the same amplitude wingbeats with much smaller muscle displacements, resulting in a sharply increasing transmission ratio (driving a decrease in *T*_*n*_) with size. Adaptations in wing hinge musculature and gearing (*2,46*) may tune the transmission ratio and enable modulation of wingstroke parameters without changing wingbeat frequency. Thus, a combination of effects on *t*_*o*_ and *T*_*n*_ likely pushes millimeter-scale fliers closer to resonance than bees, making way for extreme kinematic and morphological adaptation to facilitate maneuverability (*47*).

Our results demonstrate that the physiology of asynchronous muscle activation under cyclic strain constrains many insects to operation at or above resonance, suggestive of a widespread advantage to supra-resonant flight. The apparent ubiquity of supra-resonant flight also demonstrates that resonance tuning is not necessary for insect flight systems. While smaller insects generally beat their wings faster and have higher resonant frequencies, competing biomechanical and physiological pressures from muscle, wing, and exoskeleton make perfectly resonant wingbeats a precarious target for selection. The capacity for supra-resonance widens the phenotypic space for successful flight, opening up the possibility of evolutionary tuning of insect resonant properties for control, efficiency, or speed (*8, 12 48, 49*).

Operation above resonance may enable increased frequency-modulation capacity in asynchronous insects via modulation of resonant properties, such as during different buzzing modes in bees, or during sensory feedback driven maneuvers in flies (*31, 50*). For example, decreasing the transmission ratio by keeping the wings retracted (such as during bumblebee defense buzzing) would drastically decrease the resonant period, and could easily result in a doubling of wingbeat frequency without any modification of muscle properties (Fig. 4b) (*51*). Such a mode of frequency modulation would not be possible if bumblebees had much faster muscle stretch activation kinetics (*t*_*o*_), because the low-*t*_*o*_ region of the parameter space has a flat relationship between emergent and natural frequency (Fig. 3a, 4b). Similar modulation capacity in synchronous insects is possible by transient neurogenic frequency changes during perturbation recovery (*8, 12 48*).

Temperature-dependent modulation of wingbeat frequency via changing stretch-activation time constants may be important in thermogenesis buzzing, or for insects that fly in cold conditions such as alpine honeybees (*52*). One unique challenge faced by insects at the size of a fruitfly and smaller is that they cannot maintain flight muscle temperatures significantly above ambient temperature. This temperature constraint may cause faster-flapping insects to have a slower *t*_*o*_ since stretch-activation timescales are known to be highly temperature-sensitive (*23, 53*). Modulation of wingbeat frequency by calcium-dependent potentiation of asynchronous muscle force may also be more effective above resonance (*54*). In general, our results suggest that the timescale of asynchronous muscle’s stretch activated dynamics (*t*_*o*_) and the timescale of the mechanics of resonant spring-wing thorax (*T*_*n*_) are independent axes by which the emergent wingbeat frequency can change over long timescales by selection, or short timescales by phenotypic plasticity (Fig. 4b).

Asynchronous flight was a key evolutionary innovation that opened the possibility for superfast wingbeat frequencies, enabling insects to miniaturize and diversify. Contrary to the longstanding hypothesis of resonance tuning, our materials testing, muscle physiology, dynamical and robophysical modeling demonstrate that many asynchronous insects flap significantly above their resonant frequency. We highlight a mechanism for asynchronous supra-resonance: a tug-of-war between intrinsic physiological timescale of asynchronous muscle and the resonant mechanics of the thorax and wings. Supra-resonance also generalizes to multiple synchronous orders, despite their wingbeat frequency being determined neurally. Thus, supra-resonance is an underappreciated and widespread property of insect flight, that underscores the importance of balancing efficiency and agility across Earth’s smallest aerial locomotors.

