## Supplementary Information for "Supra-resonant wingbeats in insects"

### 1 Methods

#### 1.1 Mechanical testing experimental setup

We used an electrodynamic shaker (The Modal Shop, 2007E) to measure the material properties of the bumblebee thorax, based on the setup described in Jankauski et al (*15*). The shaker was configured horizontally, with a threaded rod used as the shaker drive shaft. A custom 3D printed plate was threaded onto the rod, onto which we affixed a piezoelectric accelerometer (PCB Piezotronics, 352A21). At the end of the threaded rod, we mounted a custom 3D printed piece that was designed to match the curvature on the anterior aspect of the bumblebee thorax, covering the approximate area of attachment of the main downstroke muscle, the dorsal longitudinal muscle (DLM). Co-linear with the shaker drive shaft, we placed a force transducer (PCB Piezotronics, 209C11). A $\#2$ threaded rod inserted directly into the force transducer, and served as the mounting surface on the posterior aspect of the bumblebee thorax, as its cross-sectional area was slightly smaller than the posterior area of attachment of the DLM. The force transducer was mounted on a three-axis stage that allowed precise positioning of the $\#2$ threaded rod on the specimen. We used a signal conditioner (PCB Piezotronics, 482C15) to power the accelerometer and force transducer, and a signal analyzer (DataPhysics, ABACUS 901) as our data acquisition system.

#### 1.2 Insect preparation and mounting

Bumblebee hives were obtained from BioBest. Individual bumblebees were anesthetized in a refrigerator for fifteen minutes prior to experimentation. We weighed each animal, and immediately isolated its thorax by removing the head, abdomen, legs, and wings. We gently filed the hairs off of the dorsal aspect of the thorax to ensure that smooth exoskeletal surfaces were free to glue to the shaker mounts. The insect was glued to the anterior mount using cyanoacrylate, which was then secured to the shaker drive shaft. The rest length of the thorax was then determined by slowly bringing the posterior mounting screw closer to the thorax using the three-axis stage. The length at which a non-zero force reading was measured on an oscilloscope was recorded as the rest length. A small drop of glue was applied to the posterior mounting screw, which was then carefully brought in contact with the thorax using the three-axis stage, compressing slightly so that the glue could set. After five minutes, the thorax was returned to its rest length.

#### 1.3 Thorax data collection and analysis

We prescribed a swept sine displacement oscillation with a $2\%$ peak-to-peak amplitude, measured with respect to a caliper measurement of the thorax length before mounting. The stimulus swept from 10 to 200 Hz, encompassing the range of measured wingbeat frequencies for *Bombus impatiens*. The sweep was conducted with a logarithmic sweep rate of 2 decades per minute.

Raw acceleration and force time series data were exported for further analysis in MATLAB. The acceleration data were high-pass filtered with a cutoff frequency of 5 Hz and integrated twice to calculate displacement. A fast Fourier transform (FFT) was applied to the displacement and force data. Then, we computed a complex stiffness (*55–57*) by dividing the transformed force, $F(i\omega)$ and displacement, $X(i\omega)$:

$$H(i\omega) = \frac{F(i\omega)}{X(i\omega)} \tag{4}$$

The gain and phase of the complex stiffness represent the stiffness and structural damping properties of a material, and can be used to compute its material properties in the context of a material model (see Supplemental Results 2.1, Fig. S1). The linear thorax stiffness we use in our modeling (see sections below) was taken to be the gain of the complex stiffness evaluated at the insect's wingbeat frequency.

#### 1.4 Muscle data collection and analysis

For delayed stretch activation (dSA) characterization, bumblebee thoraces were prepared the same way as for materials testing, with a few additional steps. The propodeum, the plate of exoskeleton overlying the posterior phragma, was carefully removed and the scutum was glued to a custom 3D-printed mount. With the phragma exposed, caliper measurements were taken of the length, width, and height of the pair of DLMs. Length was taken to be the length of the ventral-most fibers. The 3D-printed mount was screwed onto the drive shaft of the shaker, and the posterior phragma was glued to the lever arm of a dual-mode muscle motor via a tungsten rod coupler (Aurora Scientific 305C). The muscle motor was used in length-control mode as a high-resolution force sensor, and mounted on a micrometer stage allowing us to precisely control the muscle length. Muscle length was set to the zero-force length, and a saline drip from above was activated to ensure the muscle remained properly hydrated throughout the experiment. Finally, a pair of tungsten-tipped silver wire electrodes were inserted into the right DLM, one towards each end of the muscle.

A custom Simulink controller was used to prescribe stretch-hold-release-hold length trajectories to the bumblebee DLMs. The controller consisted of a Proportional, Integral, Derivative (PID) control module that was tuned manually to minimize the rise time of a representative step in length. The controller took real-time input from a fiber-optic displacement sensor (Philtec D21) that was calibrated before each experiment. We prescribed $1\%$ length changes while tetanically stimulating the muscle at 150 Hz via the implanted electrodes. Electrical stimuli were initiated 0.1 s before the onset of stretch, to allow for maximal activation of the muscle. After each trial, we increased the length of the muscle via the micrometer stage by 0.05 mm. We continued this process until the glue attaching the posterior phragma to the muscle motor broke, which usually occurred at high levels of passive tension in the muscle. We did not explicitly control for fatigue at long muscle lengths, but we stretched the muscle far beyond the $1 - 3\%$ strains that would be experienced *in-vivo*. Physiologically-relevant data was collected only within the first few trials, making it unlikely that we are significantly underestimating length-dependency of stretch activation.

Raw data was loaded into MATLAB for processing. We computed passive tension as half the resting tension in the muscle before the onset of either electrical or mechanical stimulus (accounting for the fact that we only stimulated one of the two DLMs). We extracted $t_o$ by using the *findchangepts* function (Signal Processing Toolbox, Mathworks) to identify the change in slope associated with the juncture of phases 2 and 3 of the generalized dSA response. The difference in time between this point and the time at which maximum dSA force was measured was considered to be $t_o$. We fit exponential curves to phases 3 and 4 of the dSA response to identify $r_3$ and $r_4$ using the following form and the MATLAB *fit* function:

$$g_{dSA}(t) = K_3(1 - e^{-r_3 t}) + K_4 e^{-r_4 t} \quad (5)$$

An analogous procedure was applied to the dSD response following the returning of muscle to its starting length. We pooled the measured $t_o$ and exponential rate values across dSA and dSD responses, but only values for trials with a good double-exponential fit ($r^2 > 0.9$) were used for analysis and modeling. Such filtering accounts for trials for which there was poor muscle activation, trials conducted at lengths too low for a dSA or dSD response to be noticeable, and issues related to poor fitting.

#### 1.5 Dynamically scaled robophysical flapper

We use the same dynamically-scaled robophysical flapping spring-wing system as in previous studies (*10, 20, 30, 49*). Importantly, most dynamically scaled flappers prescribe kinematics to measure aerodynamic forces (*58, 59*). This model, prescribes the forcing function (either synchronous or asynchronous) and measures the emergent kinematics. Empirically measured damped and undamped resonance frequencies were 3.50 Hz and 3.56 Hz respectively. Details of the experimental apparatus are described elsewhere, so we only provide a brief summary here.

The robophysical flapper consisted of a silicone torsion spring attached to a drive shaft with a thrust bearing and radial air bearings. An optical rotary encoder (4096 CPR, US Digital) measured motor angular position. At the bottom of the shaft was a rigid acrylic wing submerged in a tank of water. A brushless DC motor (D6374 150 KV, ODrive Robotics) was used in concert with a closed-loop motor driver enabled direct torque control at 10 kHz. The asynchronous drive was prescribed through a Simulink Desktop Real Time model, implementing the dSA transfer function (see Supplementary Methods section 1.6.2, Eq. 16) at 1 kHZ. This transfer function takes wing position from the encoder as an input and outputs the dSA force, which is multiplied by $\mu$ and is directly translated to a torque command by the motor driver. This control scheme is particularly effective as it does not require any numerical integration of the dSA equations at each time step (Eqs. 2-3).

#### 1.6 Bumblebee modeling

##### 1.6.1 Time-periodic spring-wing equation

We use the following equation to model a flapping bumblebee without taking into account stretch-activation. The calculation of each parameter in this equation is outlined below:

$$I\ddot{\phi} + \Gamma|\dot{\phi}|\dot{\phi} + \frac{k_l}{T^2}\phi = \frac{F_o}{T}\sin(2\pi f_s t) \tag{6}$$

**Wing Inertia** (**$I$**) Wing inertia is computed by the following equation (*60*), taking into account the inertia of the wings themselves as well as the added mass of air around them ($\nu$):

$$I = m_w R^2 \hat{r}_{2,m}^2 + \nu R^2 \hat{r}_{2,\nu}^2 \tag{7}$$

**Aerodynamic damping** (**$\Gamma$**) The lumped aerodynamic damping parameter is computed with the following equation (*61*), using a constant, wingstroke-averaged coefficient of drag.

$$\Gamma = \frac{1}{2}\rho C_D A_w \hat{r}_{2,s}^2 R^2 l_{cp} \tag{8}$$

**Linear thorax stiffness** (**$k_l$**) Linear thorax stiffness is a combination of stiffness from exoskeleton and active muscle. Exoskeletal stiffness was taken from the current results (Fig. S1). We defined the stiffness of the thorax in the direction of the line of action of the DLMs at free-flight wingbeat frequency (180 Hz). To get a conservative estimate of active muscle stiffness, we add twice the stiffness of one pair of flight muscles, modeling the stiffness contributions of the DLMs and DVMs which are in parallel with the exoskeleton. This is likely an overestimate because the DVMs are typically smaller than the DLMs. We use the Young's modulus of bumblebee muscle from Josephson et al. 1997, and convert it to linear stiffness using the measured dimensions of our bumblebees (*62*):

$$k_l = k_{exo} + k_{muscle} = k_{exo} + E_{muscle}(\frac{A}{L_r}) \tag{9}$$

**Transmission ratio** (**$T$**) The transmission ratio is a critical parameter of the spring-wing equation, and must be calculated carefully to ensure it is representative of conditions during free-flight. Nominally, it can be defined as the ratio between peak-to-peak wingstroke amplitude in radians and peak-to-peak muscle displacement in meters. The former is straightforward, and taken from Buchwald et al. 2010 (*7*). Muscle displacements are more challenging to estimate, given that measurements have only been taken during tethered flight of other species of bumblebee (*B. terrestris* and *B. lucorum*) (*27, 62*). While similar to *B. impatiens*, body size differences confound direct application of displacements from these species to our work. As such, we use *strain* measurements from the tethered flight literature ($\varepsilon_{tethered}$) (*27, 62*) and modify them to account for differences in tethered ($\phi_{tethered}$) and free-flight ($\phi_{free}$) stroke amplitude.

We then use these modified strains to compute muscle displacements based on measured resting muscle lengths from the bumblebees used in the current study. Both available studies on tethered bumblebee muscle strains result in nearly identical $T$ estimates using this method.

$$T = \frac{\phi_{free}}{\left(L_r \frac{\phi_{free}\varepsilon_{tethered}}{\phi_{tethered}}\right)} \tag{10}$$

**Neural driving frequency** ($f_s$) Bumblebee wingbeat frequency is defined as $f_s$, assuming that the muscle forcing to drive wingbeats is set by the nervous system. We sweep over a range of $f_s$ values to obtain the frequency response in Fig. 2b.

We are mainly concerned with the undamped resonance frequency, which is only a function of $k_l$, $T$, and $I$:

$$f_n = \sqrt{\frac{k_l}{T^2 I}} \tag{11}$$

While the damped (i.e. displacement) resonance is sometimes used in reference to insect flight, this frequency is always lower than the undamped frequency. As the displacement resonance frequency does not have a simple closed form for an oscillator with nonlinear aerodynamic damping, we simply focus on the undamped resonance frequency and note that our conclusions with respect to undamped resonance also apply to damped resonance.

**Muscle force amplitude** ($F_o$) Muscle force amplitude is set such that steady-state wingstroke amplitudes at free-flight frequency match observed free-flight amplitudes. The value of $F_o$ that achieves this in Fig. 2b is 0.65 N. Setting $F_o$ in this way ensures our model always recapitulates free-flight kinematics.

##### 1.6.2 Asynchronous spring-wing equations

We adapt a recent model of stretch-activated actuation of a robotic flapping wing for a bumblebee (*30*). The system dynamics for the sweep angle ($\phi$) of a flapping insect can be described by the following differential equation:

$$I\ddot{\phi} + \Gamma|\dot{\phi}|\dot{\phi} + \frac{k_l}{T^2}\phi = \frac{\mu}{T} f_{dSA} \tag{12}$$

Note that the left-hand-side is the same as in Eq. 6. On the right-hand-side is the non-dimensional muscle forcing, $f_{dSA}$, which is scaled by a factor $\mu$ and converted to a torque by the transmission ratio, $T$. The dynamics of $f_{dSA}$ are described by the following equation:

$$\ddot{f}_{dSA} + \alpha_2 \dot{f}_{dSA} + \alpha_3 f_{dSA} = -\alpha_3 \dot{\phi} \tag{13}$$

Here, we use the constants $\alpha_2 = \frac{\ln(1/\kappa)}{t_o}$ and $\alpha_3 = \frac{\kappa \ln^2(1/\kappa)}{t_o^2(1-\kappa)^2}$ to simplify notation. In this equation $L$ is the length of the flight muscle, $t_o$ is the timescale of dSA force rise, and $\kappa$ is the ratio of the rate of force decay to the rate of force rise ($\kappa = r_4/r_3$).

The quantities $t_o$ and $\kappa$ can be directly related to the rates $r_3$ and $r_4$ often reported in studies of asynchronous flight muscle. Curve-fitting for $r_3$ and $r_4$ can be highly sensitive to the precise form of the fit equations used, and makes it difficult to compare our muscle data to the many papers that have used slightly different fit functions or have not reported their fitting procedure in detail. Thus, we use $t_o$ and $\kappa$ to parameterize our dSA equations.

Taking the derivative of the dSA impulse response Eq. 5, setting it equal to zero, and solving for $t$, we arrive at the following relation between $t_o$, $r_3$, and $r_4$:

$$t_o = \frac{\ln(r_3/r_4)}{r_3 - r_4} \tag{14}$$

Defining the ratio $\kappa = \frac{r_4}{r_3}$, we can re-write Eq. 14 as:

$$t_o = \frac{\ln(1/\kappa)}{r_3(1-\kappa)} \tag{15}$$

From Eq. 15, $t_o$ and $\kappa$ can be calculated from $r_3$ and $r_4$. The overall dynamics are insensitive to small variation in $\kappa$ (*20*) making this formulation easier to translate across studies.

We simulate equations 2 and 3 as a system of coupled differential equations using MATLAB *ode45* and focus on the emergent wingbeat frequency at steady state. The free parameter $\mu$ is analogous to $F_o$ in Eq. 6, and is set to match emergent wingstroke amplitudes to free-flight conditions. This tuning is conducted at each grid point in Fig. 3a, so that the entire plot has a constant amplitude matching free-flight.

To compute the phase of the dSA force ($\delta$) with respect to strain, we take the Laplace transform of Eq. 13, and rearrange to derive the transfer function that relates wing angle $\Phi$ to dSA force $F_{dSA}$:

$$\frac{F_{dSA}(s)}{\Phi(s)} = \frac{-\alpha_3 s}{s^2 + \alpha_2 s + \alpha_3} \tag{16}$$

We then make the substitution $s = i\omega$ and take the complex angle to yield the following expression for $\delta(\omega)$:

$$\delta(\omega) = \tan^{-1}\left(\frac{\omega^2 - \alpha_3}{\alpha_2 \omega}\right) \tag{17}$$

To derive an expression for the phase response of a synchronous insect, we apply the method of harmonic balance (*63*) to linearize Eq. 6. Doing so yields an equivalent viscous damper $c_{eq}$ that linearizes the quadratic aerodynamic damping term and allows us to write a closed-form expression for phase.

$$c_{eq} = \frac{3}{\pi}\Gamma\omega\phi_o \tag{18}$$

The final equation for the phase of muscle force with respect to strain for a synchronous insect (black line in Fig. 3b) is:

$$\delta(\omega) = \tan^{-1}(\frac{c_{eq}\omega}{k_l/T^2 - I\omega^2}) \tag{19}$$

#### 1.7 Muscle and mechanical properties of *M. sexta*, *B. impatiens*, and *D. melanogaster*

We used a literature search to identify muscle and mechanical parameters for bees, flies, and moths to parameterize our spring-wing models. All parameters, their associated errors, and references can be found in Tables 1-3. We describe the rationale for certain parameter choices for each species in the following subsections:

##### 1.7.1 Muscle stiffness

In general, we are concerned with the dynamic muscle stiffness of the flight muscles during cyclic contraction at *in-vivo* conditions (temperature, activation, etc). For *M. sexta* and *B. impatiens*, work-loop studies make stiffness or modulus accessible from the literature for flight-relevant strain amplitudes, making sure to account for both pairs of flight muscles when appropriate. To convert between modulus and stiffness, we use the standard relationship between $E$ and $k$, $E = kL/A$, where $A$ and $L$ are the appropriate flight muscle area and length respectively for the particular data in question.

For *D. melanogaster*, reliable work-loop data at flight-relevant conditions are not available. A recent modeling study by Pons et al. 2023 performed a detailed matching procedure allowing beetle work- loop data to be scaled to Drosophila. Figure 9C of Pons et al. 2023 shows their estimated muscular elasticity for a single DLM and DVM. Using appropriate muscle dimensions (Table 3), we estimate a modulus for Drosophila muscle of $E_{muscle} = 1.1$ MPa. This is comparable to a muscle modulus estimate derived from Glasheen et al. 2017 and Loya et al 2022, who performed stretch-hold experiments on fruitfly DLM fibers ($E_{muscle} = 0.9 - 1.1MPa$). The rapid small-amplitude stretches imposed on effectively tetanized muscle likely yield a modulus that is representative of antagonist-driven stretching during flight. All of these moduli are somewhat smaller than, but similar to the modulus of bumblebee muscle during cyclic strain ($E_{muscle} = 1.6$ MPa).

##### 1.7.2 Muscle area

Muscle area for *M. sexta* and *D. melanogaster* was taken from the literature. For D. melanogaster, we multiplied the average area of a single fiber by 12, since there are 6 fibers per DLM and two DLMs in the animal. This value was then multiplied by two again to account for both pairs of flight power muscles. *B. impatiens* area measurements were measured ourselves during muscle physiology experiments.

##### 1.7.3 Muscle length

Muscle length for *M. sexta* was taken from the literature and for *B. impatiens* was measured ourselves. For *D. melanogaster*, we assume an average fiber length of 1 mm, which is representative of multiple studies (*14, 64*).

##### 1.7.4 Wing inertia

Wing inertia for *M. sexta* was directly available from the literature. For *B. impatiens* and *D. melanogaster*, we use Eq. 7 to compute inertia using wing morphometric data from the literature (see Table XX).

##### 1.7.5 Transmission ratio

Transmission ratio for *M. sexta* was directly available from the literature, and for *B. impatiens* as computed using Eq. 10. For *D. melanogaster*, we assume a peak-to-peak muscle strain of 3%. Muscle strain measurements from tethered flight in *D. melanogaster* do not exist. The range of estimated muscle strains from multiple studies ranges from 1-3%, which is consistent with strains measured in bumblebees and other asynchronous insects, (*14,27,62,65*). We choose 3% as a conservative estimate of strain since it results in the highest possible resonant frequency, but note the importance of measuring this quantity with high presicion in *Drosophila* specifically. We then compute $T$ using the equation $T = \frac{\phi_o}{\varepsilon L}$.

##### 1.7.6 dSA timescale

D. melanogaster $t_o$ was directly available from Loya et al. 2022 (*44*).

#### 1.8 Resonance frequencies of other insects

We estimated resonance frequencies for three other asynchronous insect species from Casey et al. 2023 (*22*)(plus markers in Fig. 4c): *Xylocopa californica* (carpenter bee), *Bombus centralis* (bumblebee, somewhat larger than B. impatiens), and *Musca domestica* (house fly). Resonance frequency for the dragonfly *Aeshna grandis* was reported directly by Weis-Fogh (diamond marker in Fig. 4c) (*17*).

##### 1.8.1 Linear thorax stiffness

Exoskeletal stiffness was taken directly from Casey et al (*22*). While species-specific muscle stiffness estimates are not available for these species, we very conservatively estimate that, at most, muscle stiffness will be equal in contribution to the exoskeleton in determining overall thorax stiffness. Note that for *B. impatiens*, we have demonstrated that exoskeletal stiffness is at least threefold larger than muscle stiffness, and the three insects from Casey et al. are comparable or larger in size. We should only suspect muscle stiffness to exceed exoskeletal stiffness at the size scale of *Drosophila* and smaller. Thus, this is a conservative upper bound, and we use the one-sided error bars in Fig. 4c to show the range of resonant frequencies that correspond to muscle stiffness ranging from $k_{muscle} = 0$ to $k_{muscle} = k_{exo}$.

##### 1.8.2 Transmission ratio

Thorax length was taken directly from Casey et al. (*22*). Wingbeat amplitude was assumed to be 140 degrees peak-to-peak for all three insects. Muscle strain was assumed to be $3\%$ peak-to-peak for *M. domestica* and *B. centralis*, consistent with estimates from comparable species. *X. californica* is much larger than the other hymenopterans in this study, and muscle strain tends to increase with body size (decrease with wingbeat frequency). Strain measurements using laser vibrometry of *Xylocopa* during flight show scutum displacements in the range of 200-300 um, corresponding to 3-$4\%$ strain for a bee of average thorax diameter (*66*). Thus, we conservatively estimate its strain as $4\%$, giving it a smaller transmission ratio compared to *Bombus* commensurate with its larger body size and lower wingbeat frequency.

##### 1.8.3 Wing inertia

Wing inertia was computed using Eq. 7 for all three species. Wing mass was estimated as 0.4$\%$ of body mass reported by Casey et al. (*22*). This value is representative of multiple bee species (*60, 67*). Wing mass for *M. domestica* is not directly available, however $0.4\%$ is intermediate to the approximately $0.2\%$ in fruitflies (*43*) and $> 0.6\%$ documented in larger Diptera (*60*). This figure is reasonable given that *M. domestica* is at least an order of magnitude larger in body size than *D. melanogaster*. We note that even using a fruitfly wing mass ratio does not affect our conclusion of supra-resonant flight.

Nondimensional radii of the second moment of mass were assumed to be constant among members of the same order, consistent with the rough geometric similarity of the wings of bees and flies respectively. Wing lengths were available in the literature (*67–69*). Finally, added mass inertia was conservatively estimated as a fraction of wing inertia using *B. impatiens* as a reference for Hymenoptera and *D. melanogaster* as a reference for Diptera. Added mass inertia becomes increasingly important at small body sizes, thus this approximation is conservative for the three insects in question, given the small relative sizes of *B. impatiens* and *D. melanogaster*.

##### 1.8.4 Resonance frequency and wingbeat frequency

Resonance frequency was computed using main text Eq. 1. Free-flight wingbeat frequencies were taken directly from Casey et al (*22*).

##### 1.9 Resonance frequencies from Machin and Pringle, 1959

We estimate emergent and resonant frequencies from the experiments described in Machin and Pringle, 1959 (*18*). We use the data from Figures 9 and 10. These plots show emergent (asynchronous) wingbeat frequencies from beetles as a function of virtual mass and stiffness, which are modulated using analog electronics. Resonance frequency can be computed from the given mass and stiffness using Eq. 1. In these experiments, mass and stiffness were prescribed and varied using analog electronics, not measured from real insects. Thus, the points in main text Fig. 4c from Machin and Pringle (squares) should not be interpreted as *in-vivo* conditions. Instead, these points illustrate how a totally distinct experimental methodology demonstrates supra-resonance in asynchronous flight muscle.

### 2 Extended results

##### 2.1 Frequency response of the bumblebee thorax

Further analysis of the thoracic frequency response reveals that the bumblebee thorax is compatible with a simple model of aerodynamic resonance. Thorax stiffness (gain of the complex stiffness) and structural damping coefficient (phase of the complex stiffness) were invariant with deformation frequency up to 200 Hz, marking the upper range of flight wingbeat frequencies in bumblebees. Constant material properties with frequency are the hallmark of structural damping, a material model that has been used to explain the behavior of cockroach and moth exoskeleton (*55–57, 70*). Structural damping dissipates the same amount of energy for a given displacement regardless of deformation speed, contrasting with the viscous damping model commonly used to describe many biomaterials (*71*). However, neither form of damping affects the location of a system's undamped resonant peak, indicating that we can neglect thoracic dissipation when evaluating undamped resonance in the bumblebee (*11*). Stiffness at 180 Hz (wingbeat frequency) was 4.082 $\mathrm{kN/m}$ (95$\%$ CI of the mean [3.206 4.958]) and the nondimensional structural damping constant was determined to be 0.355 (95$\%$ CI of the mean [0.204 0.507]). While weak stiffness nonlinearities have been described in some insect thoraces, time-resolved muscle strain measurements in bumblebees reveal near-sinusoidal strain trajectories in tethered flight (*62*). Additionally, under realistic deformation amplitudes ( 50 micron displacements), other hymenopteran thoraces behave remarkably linear stiffness (*15, 22*), making nonlinear thorax elasticity negligible for our global resonant frequency estimate.

##### 2.2 Dynamics of supra-resonant stretch-activated wingbeats

In this section, we describe the mechanism by which the asynchronous resonant mechanics model (main text Eqs. 2-3) produces emergent oscillatory dynamics. Supra-resonant wingbeats emerge due to a Hopf bifurcation in the spring-wing dynamics, causing oscillatory behavior above a critical threshold ($t_o^*$), at which point a pair of complex conjugate eigenvalues of the

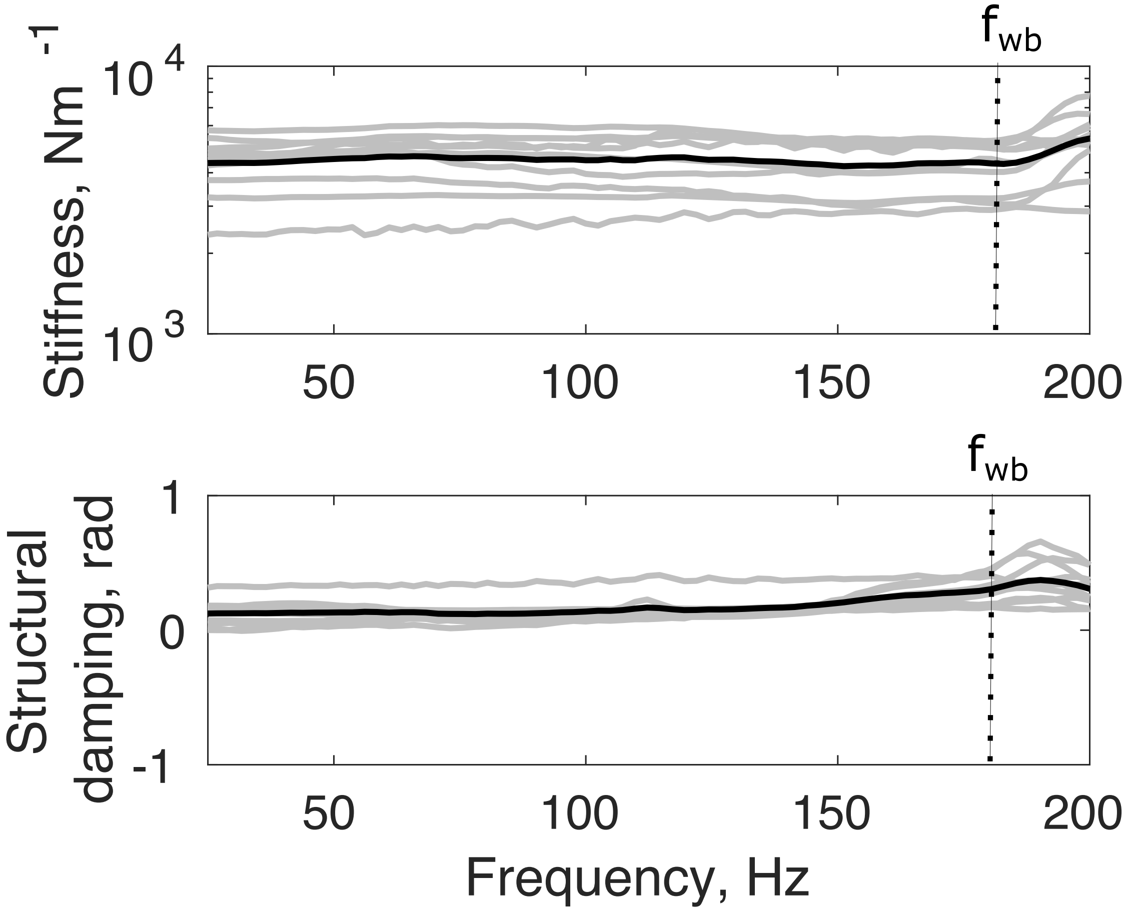


Fig. S1: Stiffness and structural damping factor as a function of deformation frequency in the bumblebee thorax. Free-flight wingbeat frequency $f_{wb}$ is annotated by the dotted lines. Solid grey lines denote individuals and solid black lines are the mean linearized system become unstable. Below $t_o^*$, the asynchronous force does not persist long enough after stretch to drive oscillation, thus acting as a dissipative element (*20,30*). $t_o^*$ depends inversely on the dSA strength $\mu$ (*20*):

$$t_o^* \approx 1.7 \frac{2\kappa(1+\kappa)}{\kappa\hat{\mu} + \sqrt{(\kappa\hat{\mu})^2 + 4\kappa(1+\kappa)^2\omega_n^2}} \tag{20}$$

In Eq. 20 above, we introduce $\hat{\mu} = \frac{\mu}{IT}$ for notational ease. Recall that $\mu$ represents the height of the dSA response of asynchronous muscle following rapid stretch. By scaling $\mu$ to maintain a fixed, realistic bumblebee wingbeat amplitude, the bifurcation critical point ($t_o$ at which the dominant real eigenvalue becomes positive) shifts down, enabling arbitrarily faster supra-resonant frequencies. Perfectly resonant wingbeats at large $t_o$ arise as the dSA dynamics become extremely slow with respect to $T_n$. This reduces Eq. 3 to a force proportional to wing velocity, analogous to a negative damping force or viscosity, causing resonant self-oscillation at an amplitude limited by the system's quadratic aerodynamic nonlinearity.

Fig. S2 demonstrates the effect of tuning $\mu$ on the bifurcation critical point and consequently emergent frequency, amplitude, and phase of the asynchronous oscillations. As $\mu$ increases, frequency increases at small $t_o$, but frequency at large $t_o$ still approaches $f_n$. Amplitudes increase with $\mu$ across all $t_o$. Note that changing $\mu$ does not change the asymptotic behavior of phase as a function of $t_o$. At large $t_o$, phase approaches $\pi/2$ (resonance) and at small $t_o$, phase approaches the same maximum value regardless of $\mu$.

#### 2.3 Work-loop representation of wingbeat limit cycles

We use the force-displacement space representation of wingbeats that was popularized by Torkel Weis-Fogh to visualize the different forces acting about the wing hinge. The flight musculature, whether synchronous or asynchronous, must supply sufficient force to deform the elastic thorax, accelerate and decelerate the inertial wings, and generate aerodynamic forces for body-weight support. Plotting each of these energetic requirements as a function of wing angle illustrates how instantaneous elastic and inertial forces relate to resonance. In Fig. S3 below, we use the non-dimensional formulation of wing torques as a function of non-dimensional wing angle derived in detail by Lynch et al. 2021 and Wold et al. 2024 (*8, 10*).

At resonance (Fig. S3b), elastic and inertial forces cancel at every point in the wingstroke, resulting in no negative work (active dissipation) required by the muscle. This cancellation, in the presence of a positive aerodynamic work requirement across the entire wingstroke, is what results in a vanishing muscle storage modulus, $E' = 0$. Off resonance, elastic and inertial forces do not cancel. Their difference manifests in excess work that must be supplied by the muscle. In the supra-resonant case (Fig. S3a), this contribution must be acceleration during the first half of each half-stroke and deceleration during the second half of each half-stroke. This is reversed in the sub-resonant case (Fig. S3c). In either case, this additional energy contribution from the muscle is what results in a nonzero storage modulus. Note that the storage modulus of the

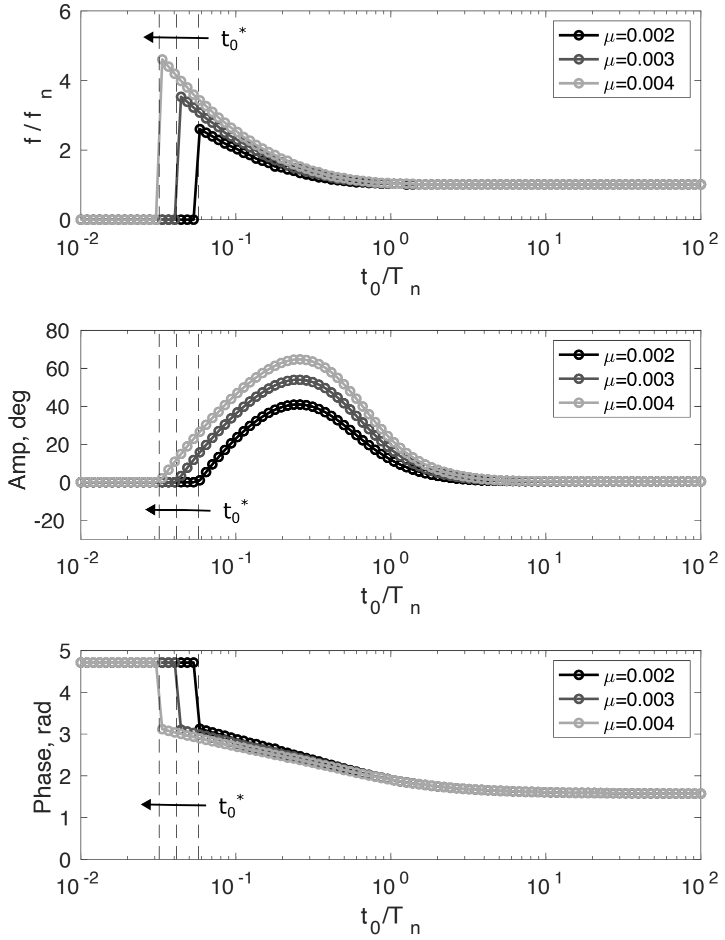


Fig. S2: Frequency (normalized by resonant frequency $f_n$), amplitude, and phase of the dSA force oscillations that emerge from simulations of an asynchronous flapping insect (Eqs. 2-3). Shades of grey show the effect of changing $\mu$ on the dynamics. Increasing $\mu$ causes a downward shift of the bifurcation critical point $t_o^*$ that enables arbitrarily supra-resonant wingbeats.

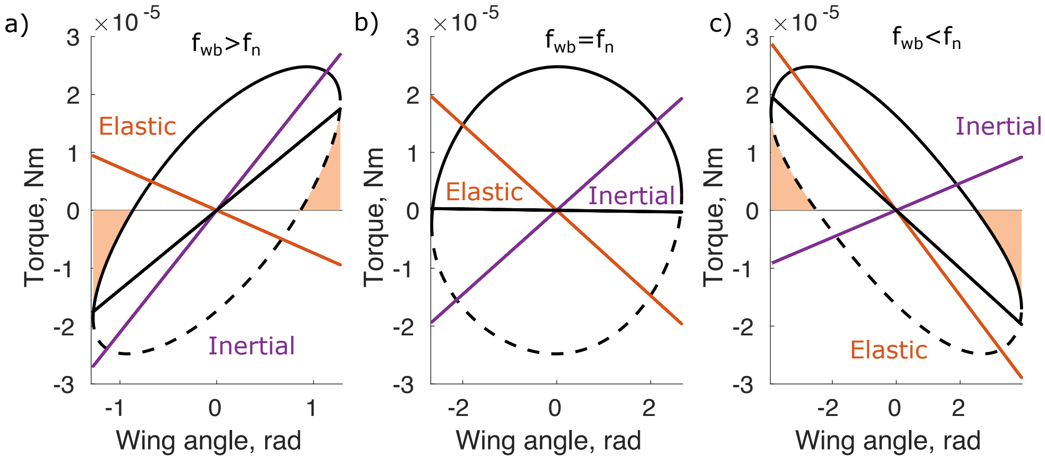


Fig. S3: Work loops at three different resonant conditions. The black ellipses show torque about the wing hinge as a function of wing angle, with the solid portion corresponding to downstroke and the dotted portions corresponding to upstroke. Shaded regions denote negative work production. Orange and purple lines show the elastic and inertial torques about the wing hinge respectively. The solid black line shows the sum of the inertial and elastic torques. a). Above resonance, inertial torques are higher in magnitude than elastic torques. This mismatch results in a positive slope (storage modulus) for the joint-level workloop. b). At resonance, elastic and inertial torques cancel exactly and the whole insect has a storage modulus of zero. Below resonance, elastic torques are in excess of inertial torques, causing the whole-insect work loop to have a negative storage modulus.

muscle reflects storage associated with the different energetic requirements of flight, and should not be associated with thoracic energy storage in isolation.

### 3 Extended discussion

#### 3.1 Series elasticity

Series compliance in the wing hinge is an important source of variation in resonant mechanics, but does not affect the emergence of supra-resonant wingbeats for the insects considered in the current work. Series elasticity in the wing hinge of insects has been demonstrated to widen the resonance curve around its global (undamped) resonance frequency (*13*). Significant series elasticity manifests in a phase difference between the deformations of the thorax and the angular movements of the wing tip. If these two timeseries are exactly in phase, the wing hinge acts as a rigid transmission. Prior work by Ando et al. 2016 in hawkmoths reveals miniscule phase lags ($\approx$ 1 ms) between the area surrounding DLM attachment and the wing-tip, such that the series-elastic phase lag is an order of magnitude less than the parallel-elastic phase lag of $\pi/2$

| Symbol | Quantity | Value | Notes |
|---|---|---|---|
| $k_{muscle}$ | Muscle stiffness | 748 N/m (*11, 29*) | For one muscle pair |
| $T$ | Transmission ratio | 2230 ± 110 1/m (*11*) | N/A |
| $I$ | Wing inertia | $5.69 \pm 0.34 \times 10^{-8}$ kg m$^2$ (*11*) | N/A |
| $f_{wb}$ | Wingbeat frequency | 25 ± 2 Hz (*11, 12*) | N/A |

Table 1: *Manduca sexta* parameters. Errors are S.E.M.

rad (10 ms for a hawkmoth wingbeat), and thus is likely negligible for moths.

Recently, Cote et al. 2025 directly measured thorax-wing phase difference in *Bombus* and found it to be 2 degrees. This is comparable to the data analyzed by Pons et al. 2022 and according to their modeling would widen the bumblebee resonance curve by less than 10 %. Thus, we can comfortably neglect series elasticity when evaluating supra-resonance in bumblebees, noting that it at most may enable some small efficiency gain compared to a completely rigid wing hinge. In bumblebees, wing hinge dynamics may play a larger role in conditions of perturbed flight or wing damage (cite Jankauski... braden + cailin preprints).

For *Drosophila*, the issue is more complex. Pons *et al.* 2022 (*13*) analyze x-ray diffraction data from Dickinson *et al.* 2005 (*72*) and suggest a more significant phase lag of approximately 22 degrees between sarcomere strain and wingbeat angle. Using muscle x-ray diffraction data to estimate has certain issues. Its time resolution is low and sarcomere strain may not be exactly in phase with bulk thorax deformation, since myofilament strain is tightly correlated with but not necessarily equal to bulk flight muscle strain (*72*). However, Pons estimates that accounting for potential variation in series elasticity, resonance peak widening of 25-50 % is likely to occur. Together with our results, *Drosophila* is supra-resonant, but only weakly and operates at close to maximal aerodynamic efficiency. However, this is likely not a general property of flies, especially those that are significantly larger than *Drosophilamelanogaster*. Evidence from blowflies (*Calliphora vicina*), which are comparable in size and wingbeat frequency to bumblebees, suggests tiny phase lags of only a couple of degrees (*73*).

| Symbol | Quantity | Value | Notes |
|---|---|---|---|
| $E_{muscle}$ | Muscle modulus | 1.66±0.25 MPa (*21*) | N/A |
| $A_{muscle}$ | Muscle area | $2.73 \pm 0.41 \times 10^{-6}$ m$^2$ | Current study |
| $L_{muscle}$ | Muscle length | $3.93 \pm 6.68 \times 10^{-5}$ mm | Current study |
| $\phi_o$ | Free-flight wingbeat amplitude | 72 ± 5 deg. (*7*) | N/A |
| $\phi_{teth}$ | Tethered wingbeat amplitude | 106 ± 7.5 deg. (*27*) | N/A |
| $\varepsilon$ | Muscle strain | 1.8 ± 0.15 % (*27*) | N/A |
| $f_{wb}$ | Wingbeat frequency | 180 ± 3 Hz (*7*) | N/A |
| $m_w$ | Wing mass | $6.88 \pm 0.34 \times 10^{-7}$ kg (*7*) | One wing pair |
| $\hat{r}_2(m)$ | N.d. second moment of wing mass | 0.44 ± 0.12 (*67*) | N/A |
| $R$ | Wing length | 0.0109 ± 0.0003 m (*7*) | N/A |
| $\hat{r}_2(\nu)$ | N.d. second moment of added mass | 0.53 ± 0.0074 (*67*) | N/A |
| $\hat{\nu}$ | N.d. added mass | 1.09 ± 0.02 (*67*) | N/A |
| $AR$ | Aspect ratio | 6.81 ± 0.0785 (*7*) | Both wing pairs |
| $t_o$ | dSA timescale | 4.4 ± 1 ms | Current study |
| $\kappa$ | dSA rate ratio | 0.17 ± 0.02 | Current study |

Table 2: *Bombus impatiens* parameters. Errors are S.E.M.

| Symbol | Quantity | Value | Notes |
|---|---|---|---|
| $E_{muscle}$ | Muscle modulus | 1.1 MPa (*44, 74*) | N/A |
| $A_{muscle}$ | Muscle area | $0.08532 \pm 0.00144$ mm$^2$ (*44*) | Pair of DLMs |
| $L_{muscle}$ | Muscle length | 1 mm , (*64*) | N/A |
| $\phi_o$ | Free-flight wingbeat amplitude | 70 ± 10 deg. (*75*) | N/A |
| $\varepsilon$ | Muscle strain | 3 % | N/A |
| $f_{wb}$ | Wingbeat frequency | 218 ± 7 Hz (*75*) | N/A |
| $\hat{h}$ | N.d. wing thickness | $5.4 \times 10^{-4}$ (*43*) | N/A |
| $\rho_{wing}$ | Wing density | 1200 kg/m$^3$ (*43*) | N/A |
| $S$ | Wing area | 3.95±0.18 mm$^2$ (*43*) | Both wings |
| $\hat{r}_2(m)$ | N.d. second moment of wing mass | 0.587 (*43*) | N/A |
| $R$ | Wing length | 2.39 ± 0.08 mm (*75*) | N/A |
| $\hat{r}_2(\nu)$ | N.d. second moment of added mass | 0.585 (*43*) | N/A |
| $\hat{\nu}$ | N.d. added mass | 1.146 (*43*) | N/A |
| $t_o$ | dSA timescale | 6.67 ± 1.8 ms (*44*) | N/A |
| $\kappa$ | dSA rate ratio | 0.02 (*44*) | N/A |

Table 3: *Drosophila melanogaster* parameters. Errors are S.E.M.

| Symbol | Quantity | Value | Notes |
|---|---|---|---|
| $k_{exo}$ | Exoskeletal stiffness | 2500 N/m (*22*) | N/A |
| $\varepsilon$ | Muscle strain | 4 % (*66*) | N/A |
| $\phi_o$ | Wingbeat amplitude | 140 deg. | N/A |
| $L_r$ | Thorax length | 7.59 mm (*22*) | N/A |
| $m_b$ | Body mass | 0.755 g (*22*) | N/A |
| $\hat{m}_w$ | Fractional wing mass | 0.4 % (*60, 67*) | Percent of $m_b$ |
| $R$ | Wing length | 15 mm (*69*) | N/A |
| $\hat{r}_2(m)$ | N.d. second moment of wing mass | 0.450 (*67*) | N/A |
| $\hat{I}_\nu$ | Added mass inertia | 1.5 | $\hat{I}_\nu = I_\nu/I_w$ |
| $f_{wb}$ | Wingbeat frequency | 120 Hz (*22*) | N/A |

Table 4: Resonance parameters for *Xylocopa californica*. Errors are S.E.M.

| Symbol | Quantity | Value | Notes |
|---|---|---|---|
| $k_{exo}$ | Exoskeletal stiffness | 400 N/m (*22*) | N/A |
| $\varepsilon$ | Muscle strain | 3 % | N/A |
| $\phi_o$ | Wingbeat amplitude | 140 deg. | N/A |
| $L_r$ | Thorax length | 2.01 mm (*22*) | N/A |
| $m_b$ | Body mass | 0.015 g (*22*) | N/A |
| $\hat{m}_w$ | Fractional wing mass | 0.4 % (*60, 67*) | Percent of $m_b$ |
| $R$ | Wing length | 5 mm (*68*) | N/A |
| $\hat{r}_2(m)$ | N.d. second moment of wing mass | 0.540 (*43*) | N/A |
| $\hat{I}_\nu$ | Added mass inertia | 1.57 | $\hat{I}_\nu = I_\nu/I_w$ |
| $f_{wb}$ | Wingbeat frequency | 160 Hz (*22*) | N/A |

Table 5: Resonance parameters for *Musca domestica*. Errors are S.E.M.

| Symbol | Quantity | Value | Notes |
|---|---|---|---|
| $k_{exo}$ | Exoskeletal stiffness | 1000 N/m (*22*) | N/A |
| $\varepsilon$ | Muscle strain | 3 % | N/A |
| $\phi_o$ | Wingbeat amplitude | 140 deg. | N/A |
| $L_r$ | Thorax length | 4.02 mm (*22*) | N/A |
| $m_b$ | Body mass | 0.13 g (*22*) | N/A |
| $\hat{m}_w$ | Fractional wing mass | 0.4 % (*60, 67*) | Percent of $m_b$ |
| $R$ | Wing length | 13 mm (*67*) | N/A |
| $\hat{r}_2(m)$ | N.d. second moment of wing mass | 0.450 (*67*) | N/A |
| $\hat{I}_\nu$ | Added mass inertia | 1.5 | $\hat{I}_\nu = I_\nu / I_w$ |
| $f_{wb}$ | Wingbeat frequency | 170 Hz (*22*) | N/A |

Table 6: Resonance parameters for *Bombus centralis*. Errors are S.E.M.

| Symbol | Quantity | Dimensions |
|---|---|---|
| $k_l$ | Linear thorax stiffness | N/m |
| $\varepsilon$ | Strain | Unitless |
| $T$ | Transmission | rad/m |
| $I$ | Wing inertia | $\mathrm{kg\ m^2}$ |
| $\Gamma$ | Aerodynamic damping parameter | $\mathrm{kg\ rad\ m^2}$ |
| $F_o$ | Muscle force amplitude | N |
| $\phi_{free}$ | Free-flight wingbeat amplitude | rad |
| $\phi_{tethered}$ | Tethered wingbeat amplitude | rad |
| $f_s$ | Wingbeat frequency | Hz |
| $R$ | Wing length | m |
| $\rho$ | Air density | $\mathrm{kg/m^3}$ |
| $A_w$ | Wing area | $\mathrm{m^2}$ |
| $C_D$ | Average drag coefficient | Unitless |
| $l_{cp}$ | Location of center-of-pressure | Unitless |
| $\hat{r}_{2,m}$ | N.d. Second moment of mass | Unitless |
| $\hat{r}_{2,s}$ | N.d. Second moment of wing shape | Unitless |
| $\hat{r}_{2,\nu}$ | N.d. Second moment of added mass | Unitless |
| $\nu$ | Added mass | kg |
| $m_w$ | Wing mass | kg |
| $L_r$ | Resting muscle length | m |
| $r_3$ | Rate of dSA force rise | $s^{-1}$ |
| $r_4$ | Rate of dSA force decay | $s^{-1}$ |
| $\mu$ | Amplitude scaling factor | Unitless |
| $\kappa$ | dSA rate ratio | Unitless |
| $t_o$ | dSA rise time | s |

Table 7: Symbol glossary
